# The Function Annotations of ST3GAL4 in Human LAMP1 and Lassa virus GP-C Interaction from the Perspective of Systems Virology

**DOI:** 10.1101/2020.04.29.068791

**Authors:** Onyeka S. Chukwudozie

## Abstract

Lassa virus (LASV) is a single-stranded RNA virus that has plagued the Sub-Saharan part of Africa, precisely Nigeria where various pathogenic strains with varied genomic isoforms have been identified. The human lysosomal-associated membrane protein 1 (LAMP1) is alternately required for the micropinocytosis of LASV. Therefore, it is of interest to understand the mechanism of action of the host LAMP1 with LASV protein during infection. The role of ST3 beta-galactoside alpha-2, 3-sialyltransferase 4 (ST3GAL4) in the interaction between LASV (glycoprotein) GP-C and the human LAMP1 is relevant in this context. Deposited curated protein sequences of both LAMP1 and LASV GP-C were retrieved for the study. The ST3GAL4 associated data was constructed and analyzed from weighted network analysis to infer the function annotations and molecular mediators that characterize the LASV infection. The gene network shows that glycoprotein sialylation, sialyltransferase enzymatic activities and glycosphingolipid biosynthesis are linked with the ST3GAL4 function. However, the physical interaction of FAM 213A, CD8B molecule and proprotein convertase subtilisin/kexin type 1 inhibitor (PCSK1N) with ST3GAL4 is intriguing in this perspective. There are eleven glycosylated asparagine sequons of the human LAMP1 but only nine were assigned a sialylated glycan cap to mediate the LASV GP-C and LAMP1 interaction having exceeded a recommended glycosylation threshold of 0.5. Therefore, the sialylated glycans of the human LAMP1 are a total of 9 and these sialylated glycans mediates the molecular recognition between LASV and LAMP1. This study therefore, predicts that there is a cellular interchange between *N-linked* glycosylation properties of the human LAMP1 and LASV glycoprotein, and sialylation functions of ST3GAL4 in LASV infectivity. Further studies and the clinical trial of this predictive model on the sialylated glycans of LAMP1 will facilitate the understanding of LASV micropinocytosis process in host cell.

## Introduction

Lassa virus (LASV) is a single-stranded RNA virus belonging to the virus family *Arenaviridae*, a family of negative-sense viruses **[1]**. Arenaviruses are enveloped viruses with a negative-sense RNA genome, consisting of two single-stranded segments named S (ca. 3.4 kb) and L (ca. 7.2 kb), each encoding two proteins with an ambisense strategy for expression **[2]**. The S segment encodes the nucleoprotein (NP) and the precursor of the envelope glycoprotein complex (GPC), while the L segment encodes the viral RNA-dependent RNA polymerase (L) and a matrix protein (Z) that is involved in virus assembly and budding **[3]**. Different strains of LASV have been identified especially in the West Africa, precisely in Nigeria. These strains are: Josiah (Sierra Leone), GA391 (Nigeria), LP (Nigeria), strain AV and few established **[4]**.

Lassa fever is a viral hemorrhagic fever caused by the Lassa virus when in contact with the causative agent Multimammate rat (*Mastomys natalensis*) **[4]**. It is reported that about 200,000 to 350,000 Lassa fever infections occur annually with approximately 5,000 deaths **[5]**. The infection, spreads from animals to humans, especially from multimammate rat and other African rat species **[4]**. This is largely the reason why the infection is difficult to curtail as most of these rat species are common in African households as adequate sanitation is not prevalent in these affected areas to reduce the rate of the spread **[4]**.

At the early stage of infection, the disease may appear asymptomatic, but at later stages the signs begin to manifest. Therefore, early clinical diagnosis and treatment is imperative for the effective eradication of the virus in the body **[6]**. During the later stage of the LASV infection especially at the incubation period of 7 to 21 days, several organs in the body gradually become dysfunctional, and the bodily manifestations of the infection include: conjunctivitis, swollen facial features, body fatigue, fever, bloody diarrhea and vomiting, nausea, constipation, cardiovascular, respiratory and nervous disorders **[6,7]**. The risk of death once infected is about one percent and frequently occurs within two weeks of the onset of the symptoms **[8]**.

Lysosomal-associated membrane protein 1 (LAMP1) also known as lysosome-associated membrane glycoprotein 1, is a protein in humans and it is encoded by the LAMP1 gene **[9]**. The human LAMP1gene is located on the long arm (q) of chromosome 13 at region 3, band 4 (13q34) **[9]**. The LAMP1 is mostly found in lysosomes, but it also located in other endosomal structures in the cell. Lysosome-associated membrane protein 1 (LAMP1) as a late endosomal co-receptor alternatively required for the internalization of LASV in a pH dependent manner **[4]**. It has been established that the intracellular LAMP1 is required for the micropinocytosis, fusion and internalization of the LASV into the cell environment **[4]**. Upon receptor binding, LASV, enters the host cell by a clathrin-independent endocytic process followed by transport to late endosomal compartments, where pH-dependent fusion of viral and cell membrane occurs **[5, 6]**. Strikingly, sodium hydrogen exchangers (NHEs) have been identified through a genome-wide small interfering RNA screen, as host factors involved in the multiplication of Lassa virus in human cells **[7]**. Based on pharmacological and genetic analysis, Iwasaki *et al*. [**14**] further validated NHE as entry factors for Lassa virus, implicating macropinocytosis in arenavirus entry, which is same for the filoviruses **[14, 15]**. The primary receptor for LASV GP is α-dystroglycan (α-DG), but it is hypothesized that the acidic pH of the late endosome destabilizes the high affinity interaction between LASV GP and α-DG, resulting in a receptor switch to LAMP1 as the alternative **[4, 16, 17]**. Lysosomal associated membrane glycoprotein acts as a secondary intracellular receptor that triggers a conformational change of the LASV that is required for the virus fusion. The proteomic interaction and stability of LAMP1 and LASV GP is regulated by the surrounding lysosomal pH conditions, where a drop in pH can destabilize LASV GP affinity for α-DG, thereby inducing potent binding to LAMP1 **[4, 17, 18]**.

The iterative haploid screens have shown that ST3 beta-galactoside alpha-2,3-sialyltransferase 4 (ST3GAL4) is imperative for the cellular internalization of LASV GP, so it is likely that mutations in the cellular functions of ST3GAL4 abrogate the ability of LAMP1 to interact with the LASV GP, which could probably be the case in *Mastomys spp* **[4]**. This signifies that a part of LAMP1 is deficient, and disrupts the endocytic process of the LASV in its major host **[4]**. After thorough literature review, this study is the first approach to exploring the relationship among ST3GAL4, LAMP1 and Lassa virus protein interactions, incorporating *in silico* methods and statistical models of a convolutional gene network. Therefore, the aim of the study is to: (I) delineate the molecular functions of ST3GAL4 using statistically derived annotations from weighted gene network analysis (II) evaluate the molecular interactions between the LASV GP-C trimeric complex and the human LAMP1, mediated by the predicted N-linked glycosylated amino residues.

## MATERIALS AND METHODS

### Protein Sequence Retrieval and Homology Modelling

Curated protein sequence of the Lassa virus glycoprotein complex (Accession number P08669) and the Human lysosome associated mediated glycoprotein 1 (Accession number P11279) were retrieved from the UniprotKB open source online repository. The *De novo* 3D protein modeling prediction of the Lassa virus polyprotein complex was generated with Raptor X, which uses a non-linear scoring function to combine homologous and structural information for a given template-sequence alignment **[20]**. Algorithmic scores and other criteria were taken into consideration before the model was considered as a good fit.

### Associated Gene Network and Annotated Functions

The Cytoscape app was used in the construction and evaluation of the associated interacting genes with ST3GAL4. Cytoscape generates hypotheses about gene function, analyzes gene lists and prioritizes genes for functional assays **[21, 22]**. Genes that also share likely functions with the query genes are predicted as well. The gene function prediction is treated as a binary classification problem. As such, each functional association network derived from the data sources are assigned a positive weight, reflecting the data sources’ usefulness in predicting the function **[23]**. A label propagation algorithm assigns a score (the discriminant value) to each node in the network. This score reflects the computed strength of association that the node has to the seed list defining the given function. This value can be used as a threshold to enable predictions of a given gene function **[23]**. A false discovery rate with an expected value (E-value) is the default statistical test used in the generation of the gene functions, because it involves multiple genomic comparison from the database. Outputs of the characterized ST3GAL4 gene ontologies and annotations were generated after apt evaluations. To decipher the functions of the gene interactions obtained from the network, the molecular function, biological process and cellular compartments were determined through the gene ontology process.

### Post Translational Modification (N-linked Glycosylation) Sites

The *N*-linked glycosylation is a very prevalent form of glycosylation and is important for the folding of many eukaryotic glycoproteins and for cell–cell and cell–extracellular matrix attachment. Therefore, the protein sequences of the GP-C of Lassa virus and the human LAMP1 were predicted for glycosylated residues that mediates their interactions. The prediction was done using NetNGlyc server **[19]**. The NetNGlyc distinguishes between glycosylated and non-glycosylated sequons. By default, predictions are only shown canonically on Asn-Xaa-Ser/Thr sequons. The Xaa signifies any amino acids asides proline due to its un-favored steric conformation. Potential scores are assigned to each position to signify it is glycosylated or not. The significant threshold is 0.5, which represents a glycosylated site. The potential scores of the glycosites are denoted by: + Potential > 0.5, ++ Potential > 0.5, +++ Potential > 0.75, and ++++ Potential > 0.90, respectively. The non-glycosylated sites are denoted by: − Potential < 0.5, −− Potential < 0.5 and −−− Potential < 0.32, respectively.

A summarized flow chart comprising the methods and analysis conducted on the virus’s protein sequences, as well as the associated data from the neural network is provided **[Figure 1]**.

**Figure 1:**
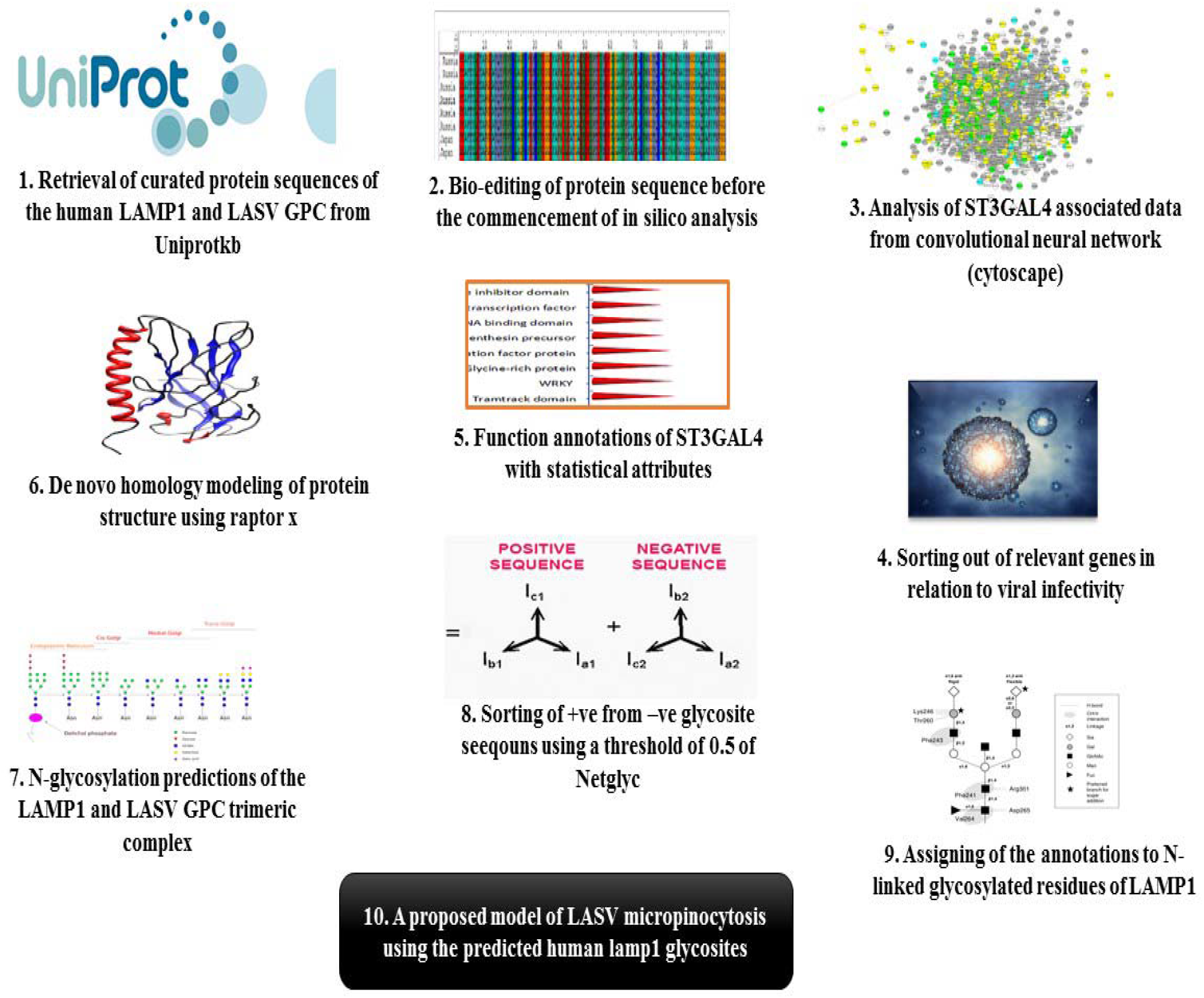
A flow chart of the summarized methods and analysis conducted.

## RESULTS

### Function annotations of ST3GAL4 from Associated Gene Network

A network visualization with annotated information was generated for ST3GAL4, which involves 20 target genes with similar expression patterns. This considered their genetic interactions, co-expression, shared pathways and similar protein structure domain. The black nodes represent 20 interacting genes and the edges are the types of interactions they share with ST3GAL4 [**Figure 2**]. The genes with absolute high interactive scores with the query gene are arranged in hierarchical regulatory order **[Table 1]**. The level of ST3GAL4 gene expression are listed in **Table 2**. The network shows that sialylation, sialyltransferase enzymatic activities and glycosphingolipid biosynthetic are linked with ST3GAL4 function. The Golgi membrane was the only gene ontology compartment generated from the network. Protein glycosylation via asparagine residue was one of the significant annotations, which was further explored in the study for the LAMP1 surface receptor and the LASV glycoprotein.

**Table 1:** Summary of genes that share similar expressions and annotations with ST3GAL4, with their respective weight scores.

| **S/N** | **Genes** | **Description** | **Co-expression** | **Physical interactions** | **Shared domain** | **Genetic interaction** |
| --- | --- | --- | --- | --- | --- | --- |
| **1** | ST8SIA5 | ST8 alpha-N-acetyl-neuraminide alpha-2,8-sialyltransferase 5 | **Y (0.03)** | **N** | **Y (0.06)** | **N** |
| **2** | ST6GALNAC6 | ST6 N-acetylgalactosaminide alpha-2,6-sialyltransferase 6 | **Y (0.03)** | **N** | **Y (0.04)** | **N** |
| **3** | ST3GAL1 | ST3 beta-galactoside alpha-2,3-sialyltransferase 1 | **Y (0.01)** | **N** | **Y (0.06)** | **N** |
| **4** | PCSK1N | Proprotein convertase subtilisin/kexin type 1 inhibitor | **N** | **Y (0.26)** | **N** | **N** |
| **5** | ST8SIA6 | ST8 alpha-N-acetyl-neuraminide alpha-2,8-sialyltransferase 6 | **N** | **N** | **Y (0.05)** | **N** |
| **6** | ST6GAL2 | ST6 beta-galactoside alpha-2,6-sialyltransferase 2 | **Y (0.02)** | **N** | **Y (0.03)** | **N** |
| **7** | ST3GAL3 | ST3 beta-galactoside alpha-2,3-sialyltransferase 3 | **N** | **N** | **Y (0.06)** | **N** |
| **8** | ST8SIA3 | ST8 alpha-N-acetyl-neuraminide alpha-2,8-sialyltransferase 3 | **N** | **N** | **Y (0.05)** | **N** |
| **9** | ST8SIA2 | ST8 alpha-N-acetyl-neuraminide alpha-2,8-sialyltransferase 2 | **N** | **N** | **Y (0.06)** | **N** |
| **10** | ST3GAL2 | ST3 beta-galactoside alpha-2,3-sialyltransferase 2 | **N** | **N** | **Y (0.06)** | **N** |
| **11** | ST3GAL6 | ST3 beta-galactoside alpha-2,3-sialyltransferase 6 | **N** | **N** | **Y (0.06)** | **Y (0.001)** |
| **12** | ST6GALNAC2 | ST6 N-acetylgalactosaminide alpha-2,6-sialyltransferase 2 | **N** | **N** | **Y (0.05)** | **Y (0.003)** |
| **13** | ST8SIA4 | ST8 alpha-N-acetylneuraminide alpha-2,8-sialyltransferase 4 | **N** | **N** | **Y (0.05)** | **N** |
| **14** | ST8SIA1 | ST8 alpha-N-acetylneuraminide alpha-2,8-sialyltransferase 1 | **N** | **N** | **Y (0.05)** | **Y (0.001)** |
| **15** | ST3GAL5 | ST3 beta-galactoside alpha-2,3-sialyltransferase 5 | **N** | **N** | **Y (0.05)** | **Y (0.001)** |
| **16** | ST6GAL1 | ST6 beta-galactoside alpha-2,6-sialyltransferase 1 | **N** | **N** | **Y (0.05)** | **Y (0.001)** |
| **17** | CD8B | CD8b molecule | **N** | **Y (0.89)** | **N** | **N** |
| **18** | FAM213A | Family with sequence similarity 213member A | **N** | **Y (0.21)** | **N** | **N** |

**Note:** Y-Yes, N-No

**Table 2:**
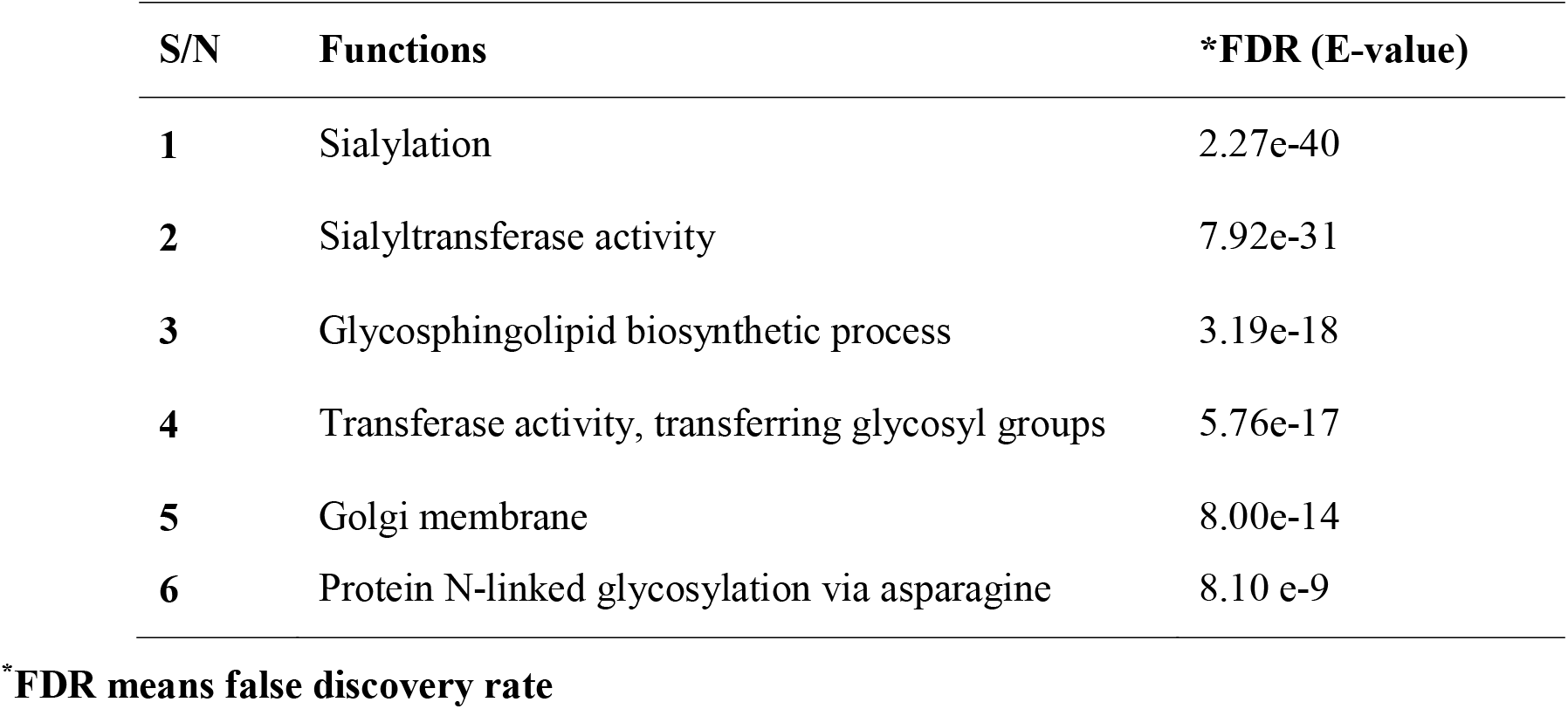
Annotations of ST3GAL4 from the Network Analysis

**Figure 2.**
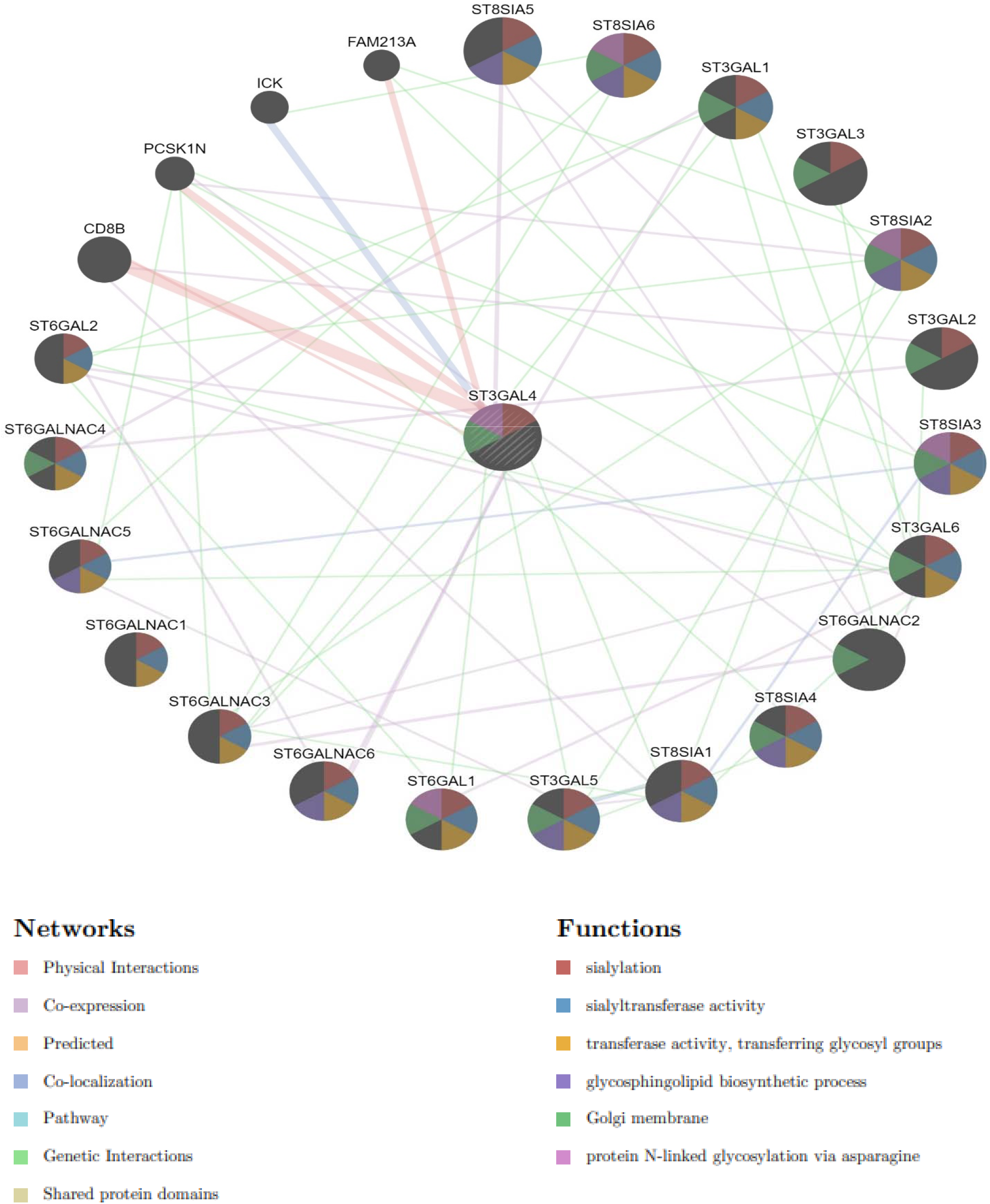
A concentric network visualization layout with annotated information for 20 important genes interacting with ST3GAL4. It also includes physical and genetic interactions, co-expression, shared pathways and protein structure domain represented by the edges between the nodes. Black nodes represent the 20 important genes, and the lines represent the different interactions of the genes. The different colour displayed in the nodes represents the annotated functions of the ST3GAL4 is and also its specific pathway types with related genes.

Out of the interactive genes that shares similar expression pathway, CB8B molecule, FAM 213A and proprotein convertase subtilisin/kexin type 1 inhibitor (PCSK1N) were the only genes that share physical interaction with ST3GAL4 with their respective interactive scores. But, CB8B interaction exhibited the highest confidence score in the network, suggesting that there could be a functional relationship both genes share in LASV infectivity.

Genes that are co-expressed with ST3GAL4 are 8, which were mostly the sialytransferase enzymatic functioning genes and those with alpha-N-acetyl-neuraminide chemical moieties. Although the co-expression interactions are transient as the weight scores are lower. These co-expressed genes could be controlled by the same transcriptional program, similar function annotations or same complex protein pathway. Genes with sialytransferase activities that share same domain with ST3GAL4 are 16 [**Figure 2**]. None of the genes are co-localized in the network meaning that the misplacement of a gene does not affect the other. The interpretations of ST3GAL4 interactions with other genes in the network are summarized in **Table 1**.

From the observed physical interactions between the triads ST3GAL4, PCSK1N and CD8B molecule, a separate network visualization was constructed where both PCSK1N and CD8B were added as the query genes to infer what other genes are recruited in the network as well as additional annotations that could characterize the ST3GAL4 and other major interactor’s role in LASV infection **[Figure 3]**. CD8B molecule annotation is enriched in immune response as it mostly recruits other T cell subsets with their functional role **[Table 3]**.

**Table 3:** Annotations of CD8B from the Network Analysis

| S/N | Functions | *FDR (E-value) |
| --- | --- | --- |
| 1 | T cell receptor complex | 3.61e-4 |
| 2 | Coreceptor activity | 1.11e-3 |
| 3 | MHC protein binding | 1.27e-3 |
| 4 | MHC class 1 protein binding | 7.48 e-2 |

\*FDR means false discovery rate

**Figure 3.**
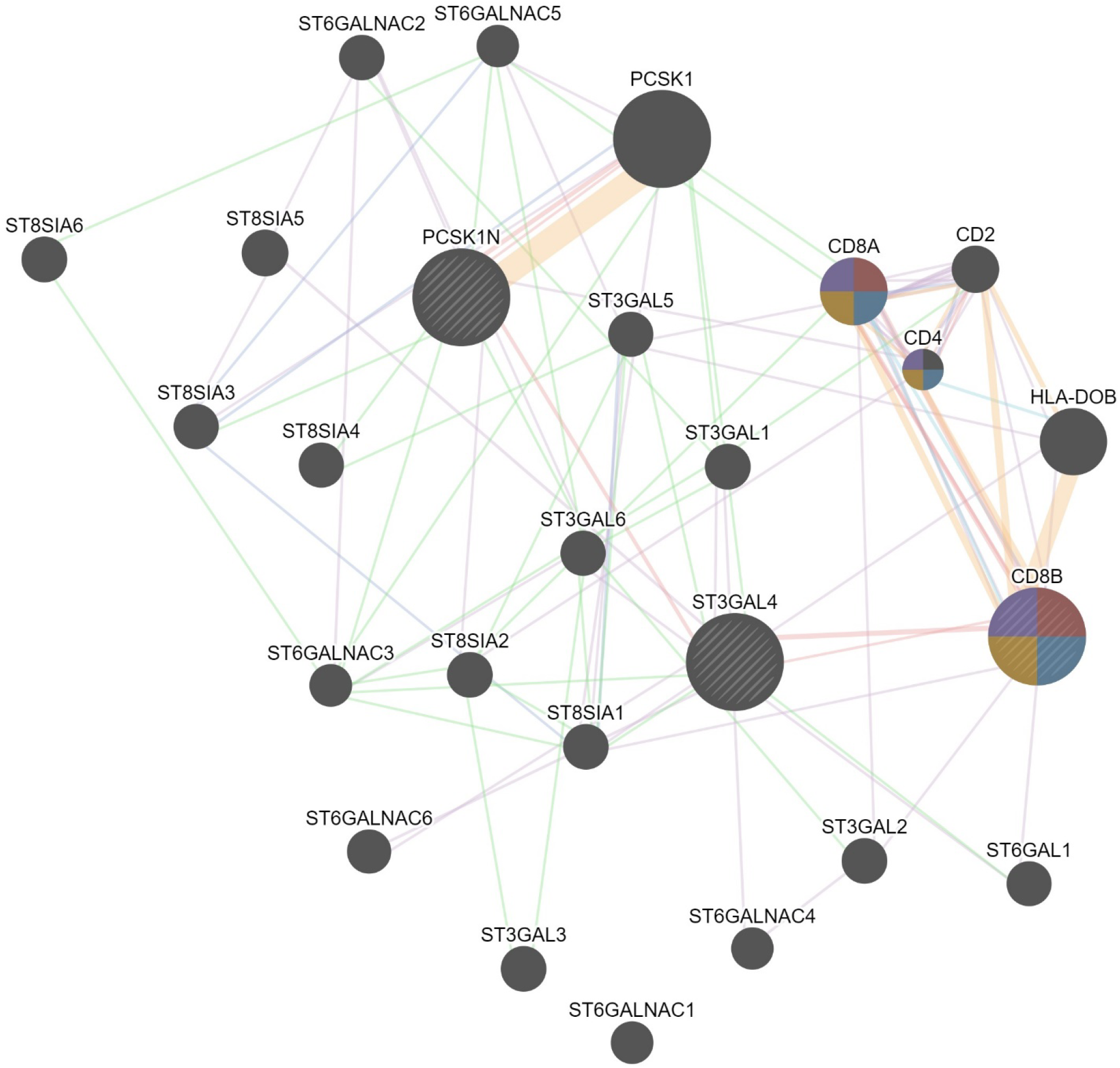
A weighted network consisting of ST3GAL4, CD8B and PCSK1N query genes connected via a strong link (physical interactions). Genes with similar expression patterns were recruited in the network.

### N-glycosylation of the GP-C of Lassa virus

From the prediction of the N-GlcNAc residues of both the GP1 and GP2 subunits of the glycoprotein, at position 89 of the asparagine sequons, it was predicted as the only negative glycosite, which was lower than the threshold value of 0.5, while other asparagine positions and sequences succeeding it were confirmed as positive glycosites, as they exceeded 0.5 **[Table 4; Figure 4]**. These glycosylated residues mediate the Lassa virus interaction with human LAMP1.

**Table 4:**
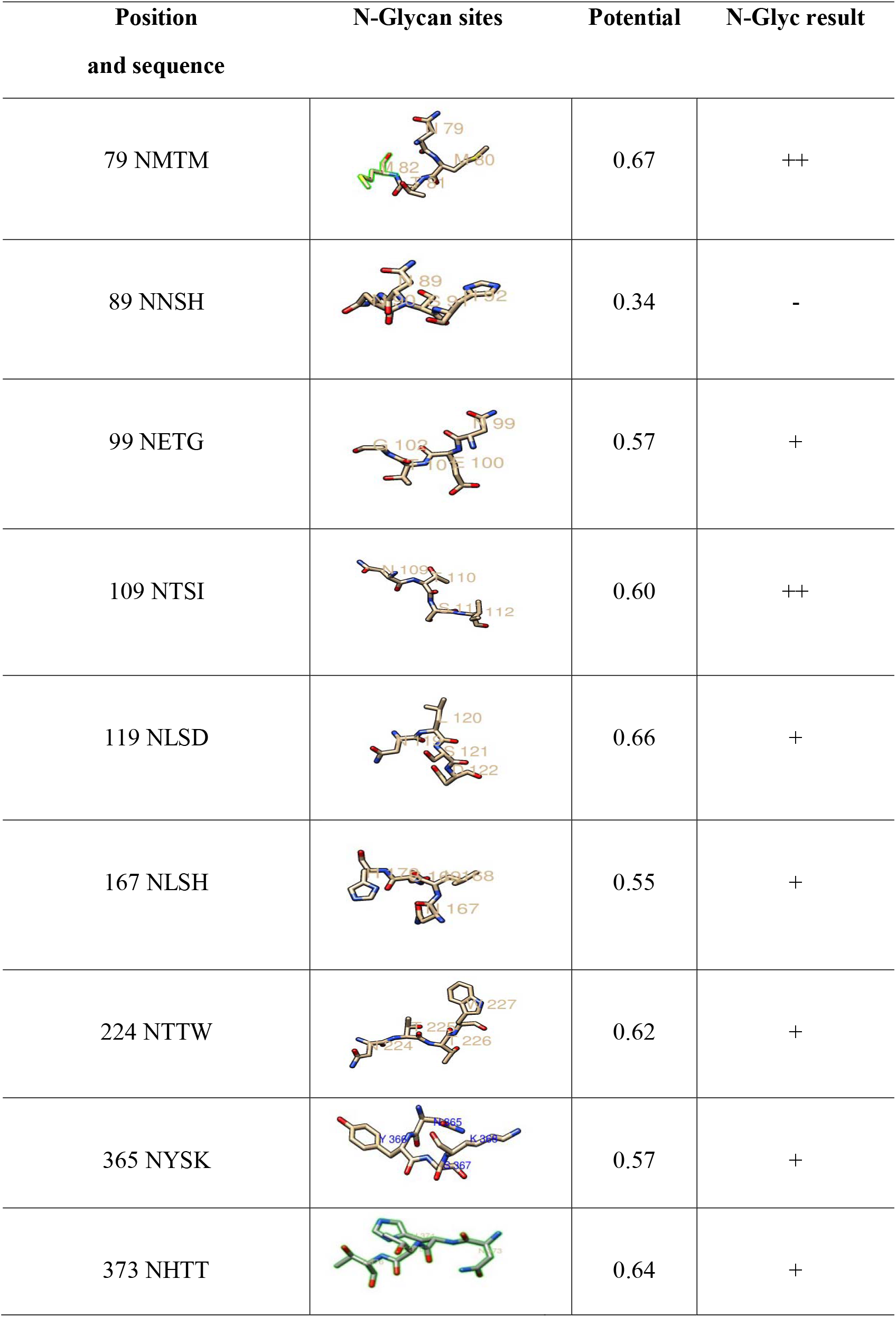
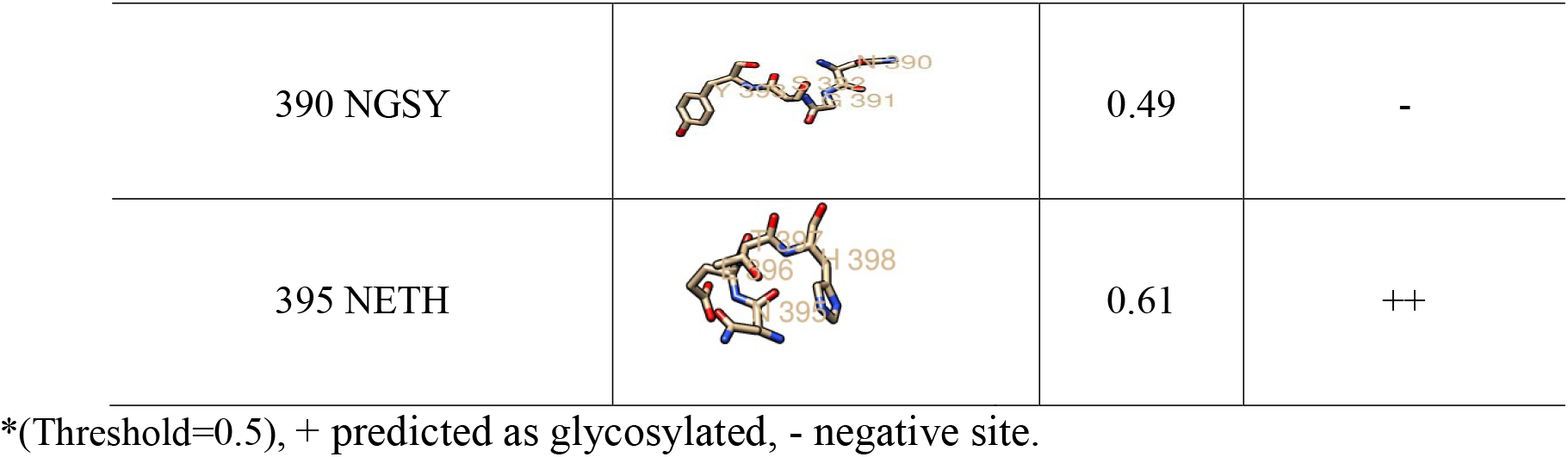
The N-glycosylated residues of Lassa virus pre-glycoprotein complex

| Position<br>and sequence | N-Glycan sites | Potential | N-Glyc result |
| --- | --- | --- | --- |
| 79 NMTM |  | 0.67 | ++ |
| 89 NNSH |  | 0.34 | - |
| 99 NETG |  | 0.57 | + |
| 109 NTSI |  | 0.60 | ++ |
| 119 NLSD |  | 0.66 | + |
| 167 NLSH |  | 0.55 | + |
| 224 NTTW |  | 0.62 | + |
| 365 NYSK |  | 0.57 | + |
| 373 NHTT |  | 0.64 | + |
| 390 NGSY |  | 0.49 | - |
| 395 NETH |  | 0.61 | ++ |

\*(Threshold=0.5), + predicted as glycosylated, - negative site.

**Figure 4:**
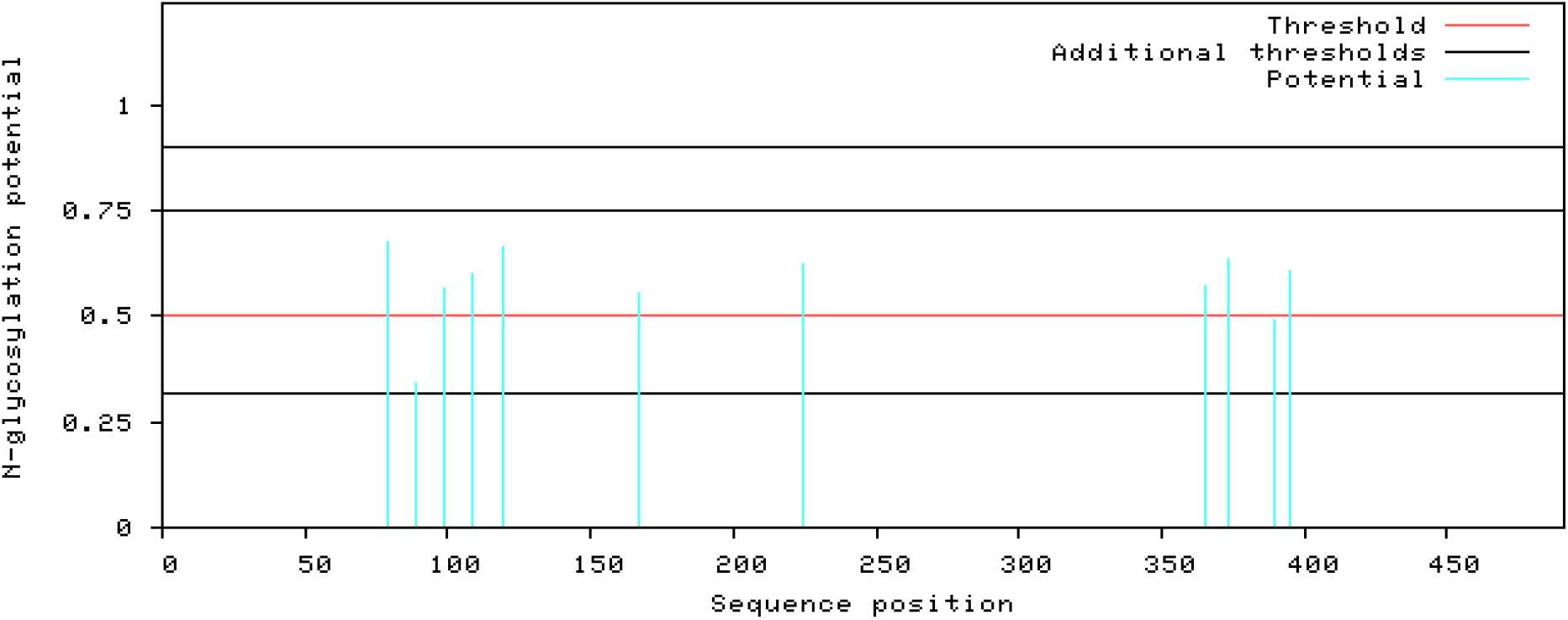
Graphical output of the thresholds and potentials of the N-glycosylated Lassa virus GP-C complex. The graph illustrates predicted N-glyc sites across the protein chain (X-axis represents protein length from N-to C-terminal). A position with a potential (vertical lines) crossing the threshold (horizontal lines at 0.5) is predicted glycosylated. All of the predicted potential residues exceeded the threshold of 0.5, as shown in the graph. The signal precursor had no glycosylated residue as N-glycosylation starts from the GP1 complex. LASV GP2 glycosylated residues starts from position 365 as graphically displayed.

### Post Translational Modification (N-glycosylation) of the Human LAMP1

The glycosylated sequons of the human LAMP1 was predicted to determine residues that mediates cellular recognition of the LASV. The glycosylated asparagine sequons are summarized in **Table 5**

**Table 5:** The N-glycosylated residues of the Human LAMP1

| Position and sequence | Potential | N-Glyc result |
| --- | --- | --- |
| 37 NGTA | 0.76 | +++ |
| 45 NFSA | 0.59 | + |
| 62 NMTF | 0.59 | + |
| 76 NRSS | 0.63 | ++ |
| 84 NTSD | 0.69 | ++ |
| 103 NFTR | 0.80 | +++ |
| 107 NATR | 0.66 | + |
| 121 NLSD | 0.71 | ++ |
| 130 NASS | 0.65 | + |
| 165 NVTV | 0.71 | ++ |
| 181 NSSF | 0.45 | - |
| 223 NVSG | 0.64 | + |
| 228 NGTC | 0.56 | + |
| 241 NLTY | 0.73 | ++ |
| 249 NTTV | 0.69 | ++ |
| 261 NKTS | 0.70 | + |
| 293 NASS | 0.53 | + |
| 322 NGSL | 0.71 | ++ |

\*(Threshold=0.5), + predicted as glycosylated, - negative site.

## SIALYLATED GLYCANS

The *N*-Acetyl neuraminic acid (Neu5Ac), and its attachment to the terminal end of LAMP1 N-glycans linked to the to the C-3 of the adjacent galactose (α 2,3 linkage) (Neu5A(α2,3)Gal(b1-4)(Fuc(α1-3))GlcNAc) of the protein sugar was structurally designed using DrawGlycan SNFG **[Figure 5]**.

**Figure 5.**
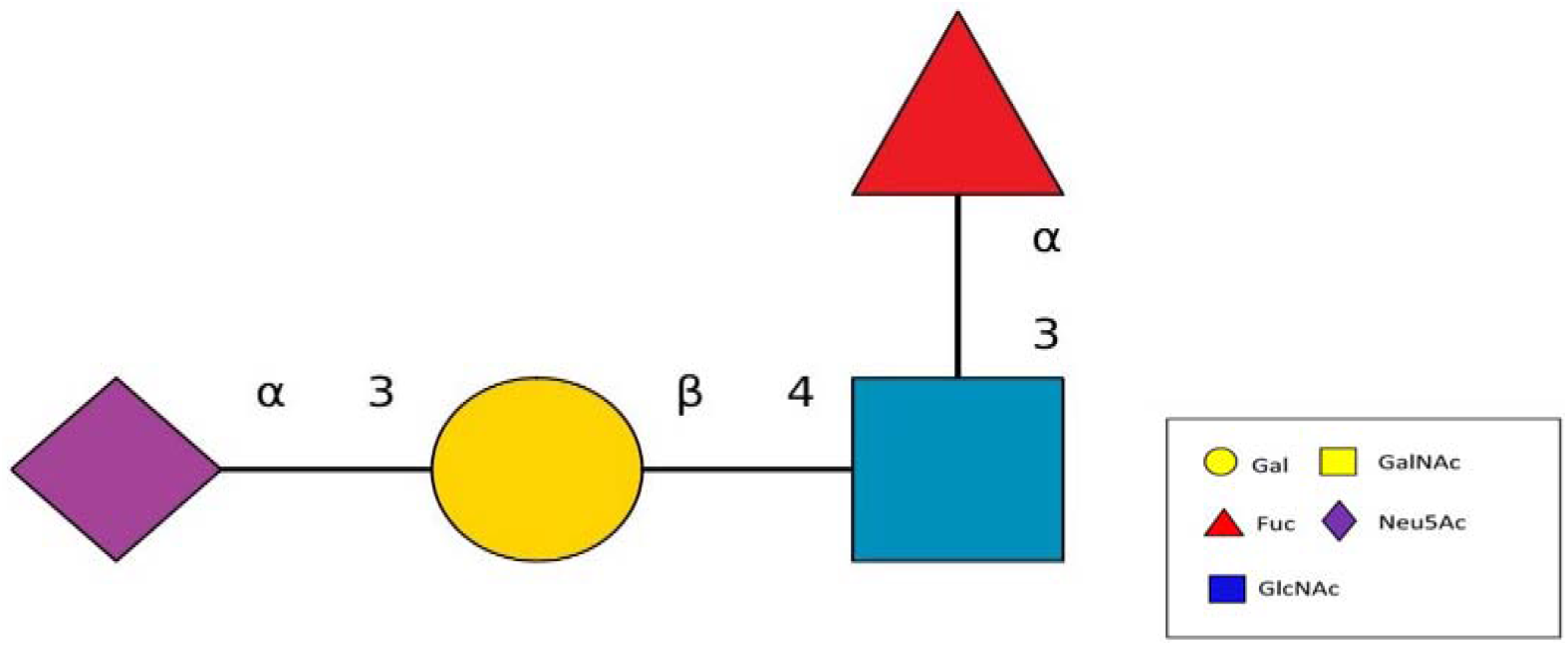
The α2,3 anomeric linkage of the Neu5Ac and the galactose sugar catalyzed by neuraminidase. The structural moiety of the α2,3 linkage Neu5Ac, which is the biologically relevant form relating to the Lassa virus infection and ST3GAL4, which is beta galactoside alpha 2,3 glycosidic bond.

### Assigning of Sialylated Glycan Cap to the Human LAMP1 Receptor

The human LAMP1 glycosylated residues whose potential score exceeds the threshold 0.5 is assumed to be highly glycosylated. Therefore, the Neu5Ac moiety attaches to the terminal end of Gal and GlcNAc of the residues with higher glycosylation which are represented with double and triple pluses (++ and +++), as the mannosylation level varies across the predicted glycosylated residues **[Table 6]**. The ++ and +++ glycosylated sequons of the human LAMP1 have a dominant level of Man9-GlcNAc2, and are consequentially capped with sialic acid moiety. The seqouns that were filtered based on their higher scores are presented in a graphical form [**Figure 6**]. From the graphical output the asparagine sequon at position 103, is comparatively higher in glycosylation than the other asparagine seqouns, which is as result of the flanking amino acids (threonine at +2 position) preceding the asparagine residue. All of these filtered seqouns were assigned sialic caps and predicted to be the human LAMP1 sialylated glycans that mediates the receptor interaction with LASV [**Figure 6**].

**Figure 6:**
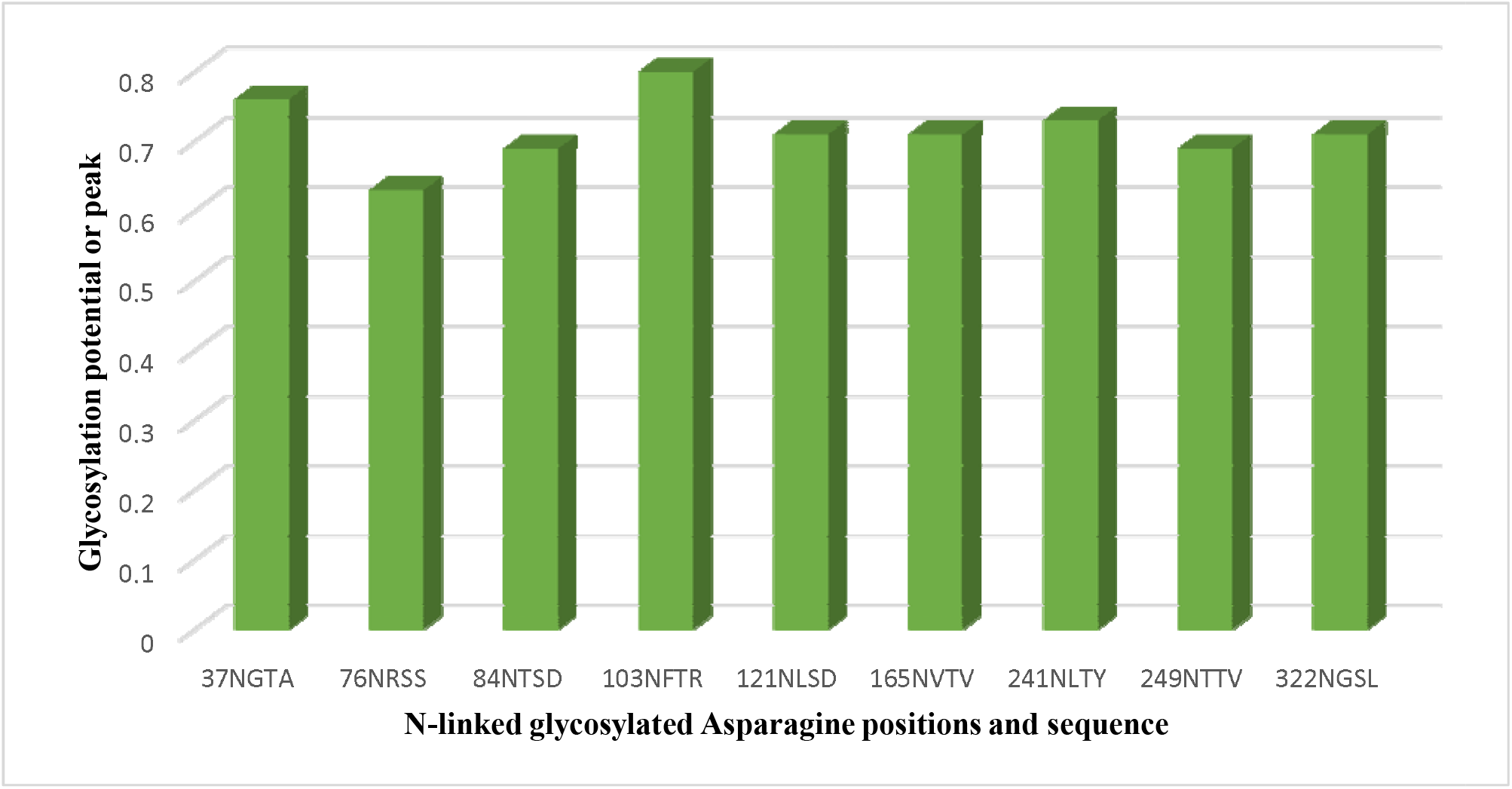
The glycosylated peak of the human LAMP1 asparagine seqouns highly dominated with high mannosylation and exceeding the threshold of 0.5.

A model was designed to represent the human LAMP1 sialylated glycans and their interaction with the Lassa GP-C **[Figure 7]**.

**Figure 7:**
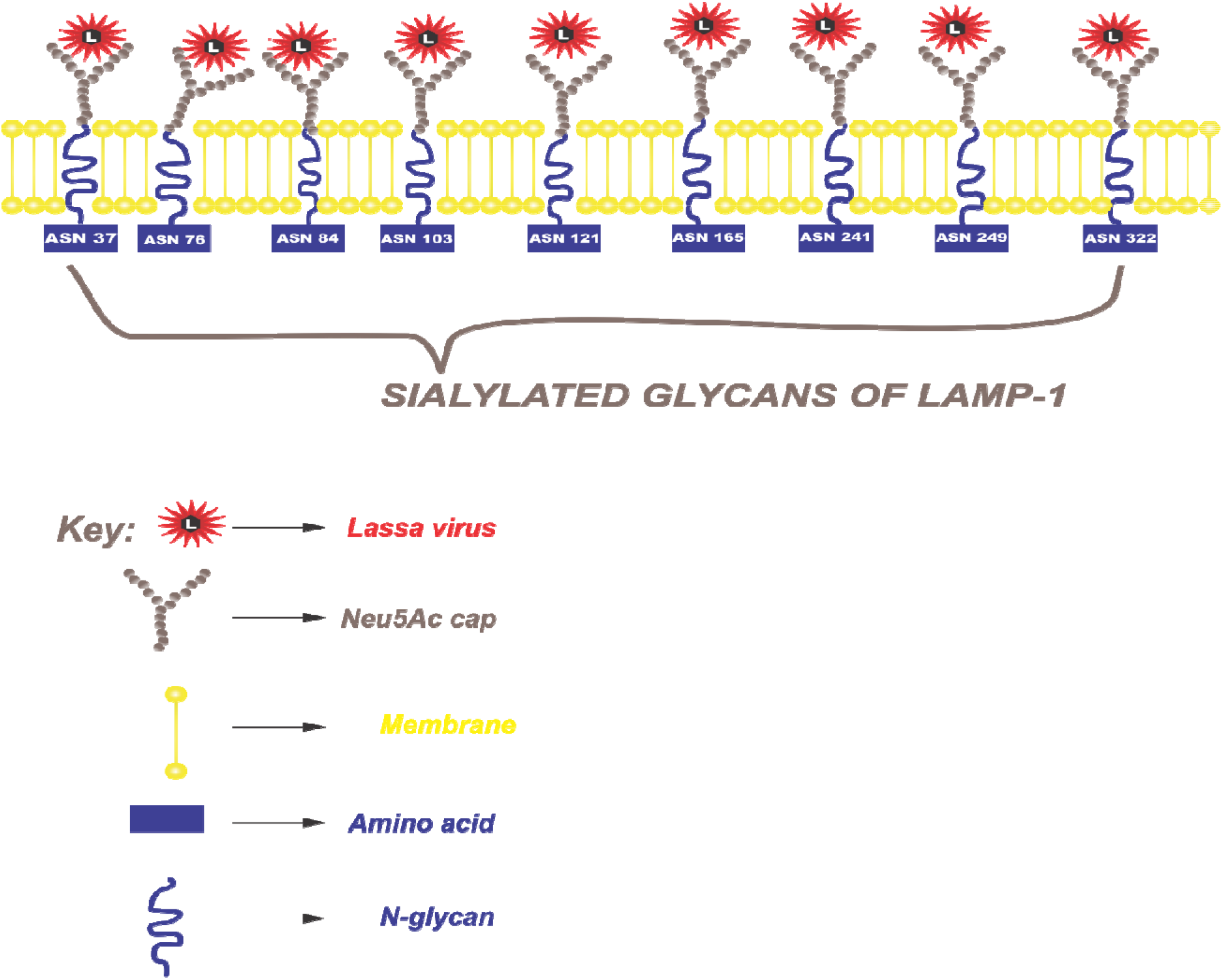
Sialylated glycans of the human LAMP1. The glycoconjugate of sialic acid and N-glycans (Neu5A(α2,3) Gal(b1-4)(Fuc(α1-3))GlcNAc) is recognized by LASV GP-C complex facilitating its cell recognition and cellular entry through the host receptor. The virus-encoded neuraminidase protein catalyzes removal of Neu5Ac from the cell surface and viral glycoproteins to release newly formed virions. The glycosylated Lassa virus protein mediates the attachment of the virus to the receptor.

## DISCUSSION

This study described the interactions involving the Lassa virus GP-C complex and endosomal receptor (LAMP1) considering specific cellular functions, such as the sialylation role of the ST3GAL4 and the N-linked glycosylation function of the human LAMP1. The analyses revealed that the human LAMP1 is highly glycosylated with most asparagine seqouns attaining and exceeding the threshold of 0.5. The asparagine seqouns with double and triple pluses were assigned sialylated glycan caps, which represented the glycans of the human LAMP1 that mediates the LASV recognition to the surface receptor. The N-glycosylation attributes of the LAMP1 contributes to its structure, cell to cell recognition and adhesion with the virus protein. From the study, it is apparent that majority of lysosomal receptors are heavily glycosylated, thereby enhancing the viral exploration of the late endosomal receptors for endocytic process. Thus, this suggests that asides the pH properties of the late endosomal receptor as widely studied, the glycosylated attributes of the endosomal receptor justify the various viral exploitations of the lysosomal proteins.

The associated data from the network analysis reveals glycoprotein sialylation is the most significant function of ST3GAL4 with its statistical attribute. Sialylation is the addition of sialic acid groups to oligosaccharides and carbohydrates as the terminal monosaccharides **[24]**. Sialic acid is the N or O substituted derivatives of neuraminic acid (9-carbon monosaccharide) and are well expressed and distributed on cellular surfaces of glycoproteins of eukaryotic cells **[24]**.

Sialic acids as terminal sugars of *N*- and *O*-glycosylated macromolecules are decisive for the functions of glycoconjugates both in soluble form and integrated into cell membranes **[24, 25]**. The sialyltransferase function is performed by ST6GALNAC3 which also belongs to the family of sialyltransferases that transfer sialic acids from CMP-sialic acid to terminal positions of carbohydrate groups in glycoproteins and glycolipids, which is the case in LAMP1. From the neural network, ST6GALNAC3 is co-expressed and shares a domain with ST3GAL4. It facilitates the transportation of the Neu5Ac, which is the activated form of NeuAc to the Golgi body. This is done by an enzyme called CMP-N-acetylneuraminate-beta-galactosamide-alpha-2,3 sialyltransferase encoded by ST3GAL4 **[25]**. By the transportation of the Neu5Ac to the Golgi body, Neu5Ac is added to the oligosaccharide chains of the LAMP1 glycoproteins. This corroborates the result obtained from the gene network that Golgi body is the cellular factory for sialylation process.

The N-glycans of LAMP1 is modified with α2,3-linked sialic acid (SA) moieties to enable signal recognition and adhesion, glycoprotein function and stability of the LAMP1 and LASV glycoprotein spike complex. The SA is installed mainly in two different ways at the terminals of N-glycans: linked to position C-6 of the adjacent galactose (α2,6 linkage type) or to C-3 of this sugar (α2,3linkage type).

An intriguing observation was the physical interaction between an alkaline serine proteinase inhibiting encoding gene, proprotein convertase subtilisin/kexin type 1 inhibitor (PCSKIN), and ST3GAL4. This interaction was the most significant in the network. Physical interactions often involve high-affinity, stable binding, which can be easily detected by biochemical methods, but may alternatively involve transient phenomena, such as protein modification or cleavage **[26]**. This clearly implicates proprotein convertase subtilisin/kexin type 1 (PCSK1) as the endopeptidase that facilitates the dissociation between the LASV trimeric complex. Therefore, PCSK1N could play a functional role in inhibiting the co-translational cleavage process mediated by PCSK1 for the protein maturation of GP1 and GP2 before molecular interactions with the sialylated glycan of LAMP1. This corroborates with the studies conducted by Sakai *et al*. **[27]** that proprotein convertase site 1 protease (SIP) was the enzyme that facilitates the cleavage between the GP-C. Numerous studies have incorporated computational pipeline to study several viral host cell interactions. For instance, a study by Chukwudozie **[15]**, revealed the interacting amino acids between the Ebola virus proteolytical processed glycoprotein and the NPC1 receptor using computational predictions. The research fully portrayed the concept of virus-host specificity using the C domain of the NPC1 by incorporating mutational studies and structural modelling.

## CONCLUSION

Lassa virus have a high affinity for sialylated glycans embedded in the endosomal glycoproteins of the host receptor. Analyzed data shows that ST3GAL4 is linked with LAMP1 during cellular recognition of LASV, proving that the role of ST3GAL4 in the protein interaction between LASV GP-C and the human LAMP1 is indispensable. These preliminary data provide insights for the design and development of therapeutics towards the eradication of LASV. A clinical approach of this study will further shed light on the multi-faceted mechanisms that govern the LASV internalization process involving sialylated glycans of the LAMP1 host receptor.

## Acknowledgement

I would like to appreciate the assistance of Treeline Innovations (research section) and its management headed by Mr. Olaolu Ibitoye for their technical assistance and other services rendered during the course of this research. I also thank Charlotte Ndiribe (Ph.D.) University of Lagos for her preliminary critics and editorials during the preparation of the manuscript. The editorial advice and research logistics rendered by Ms. Ndudi Lever, during the research and manuscript preparations is highly appreciated as well.

